# Dissociation of tic generation from tic expression during the sleep-wake cycle

**DOI:** 10.1101/2020.06.29.176974

**Authors:** Esther Vinner Harduf, Ayala Matzner, Katya Belelovsky, Izhar Bar-Gad

**Affiliations:** The Leslie & Susan Goldschmied (Gonda) Multidisciplinary Brain Research Center, Bar-Ilan University, Ramat-Gan, 52900, Israel

## Abstract

Motor tics, the hallmark of Tourette syndrome, are modulated by different behavioral and environmental factors. A major modulating factor is the sleep-wake cycle in which tics are attenuated to a large extent during sleep. This study demonstrates a similar reduction in tic expression during sleep in an animal model of chronic tic disorders and investigates the underlying neural mechanism. We recorded the neuronal activity during spontaneous sleep-wake cycles throughout continuous GABA_A_ antagonist infusion into the striatum. Analysis of video streams and concurrent kinematic assessments indicated tic reduction during sleep in both frequency and intensity. Extracellular recordings in the striatum revealed a state dependent dissociation between motor tic expression and their macro-level neural correlates (“LFP spikes”) during the sleep-wake cycle. LFP spikes, which are highly correlated with tic expression during wakefulness, persisted during tic-free sleep and did not change their properties despite the reduced behavioral expression. Local, micro-level, activity near the infusion site was time-locked to the LFP spikes during wakefulness but this locking decreased significantly during sleep. These results suggest that while LFP spikes encode motor tic generation and feasibility, the behavioral expression of tics requires local striatal neural activity entrained to the LFP spikes, leading to the propagation of the activity to downstream targets and consequently their motor expression. These findings point to a possible mechanism for the modulation of tic expression in TS patients during sleep and potentially during other behavioral states.

**Summary:** The expression of motor tics, the defining symptom of Tourette syndrome, is modulated by environmental and behavioral factors. In this study, we explored tic modulation during the sleep-wake cycle and its underlying neurophysiological mechanism, using the rat model of chronic motor tic expression. Behaviorally, during sleep, tic frequency and intensity declined considerably. Physiologically, however, the macro-level neural correlates (termed “LFP spikes”) of tics persisted throughout the sleep-wake cycle, whereas the micro-level correlates were reduced during sleep. This dissociation between neuronal activity and its behavioral expression leads not only to a better understanding of tic modulation during sleep but may also suggest potential ways of affecting tic expression during natural behavior and when exposed to external modulating factors.

**Highlights:**

- The wake-sleep cycle modulates tic expression in chronically tic-expressing rats
- Sleep attenuates tics in the rats reminiscent of the phenomenon in TS Patients
- During tic-less sleep tic generation persists via LFP spikes
- During sleep tic expression is blocked via striatal activity desynchronization

## Introduction

Tics are the defining symptom of Tourette syndrome (TS) and other tic disorders. They manifest as sudden, rapid, recurrent movements (motor tics) or sounds (vocal tics) (American Psychiatric Association, 2013). Tics vary in frequency and severity over multiple timescales (Leckman et al., 1998; Peterson and Leckman, 1998). This waxing and waning phenomenon is poorly understood but has been associated with both environmental and internal factors (Silva et al., 1995; Conelea and Woods, 2008; Caurín et al., 2014; Barnea et al., 2016; Buse et al., 2016). One key factor whose tic-attenuating effect is still a topic of heated debate, is sleep (Oksenberg, 2020). In healthy subjects, falling asleep is accompanied by muscle tone reduction (Jacobson et al., 1964; Chase, 1990). Early studies of TS patients reported an absence of tics during sleep (Shapiro et al., 1973; Rothenberger et al., 2001); however, later polysomnographic studies reported their persistence (Glaze et al., 1983; Jankovic et al., 1984; Jankovic and Rohaidy, 1987; Silvestri et al., 1990, 1995; Fish et al., 1991; Cohrs et al., 2001) and found that multiple properties of tics undergo significant changes. Comparisons to tic expression during wakefulness indicated a reduction in tic frequency and intensity during sleep (Silvestri et al., 1990, 1995; Fish et al., 1991). The tics expressed during sleep were characterized primarily as simple tics that differed from observed mixture of simple and complex tics during wakefulness (Cohrs et al., 2001).

The basal ganglia (BG) play a key role in the pathophysiology of tic disorders (Peterson et al., 2003; Bloch et al., 2005; Kalanithi et al., 2005; Worbe et al., 2015; Ramkiran et al., 2019), particularly through the reduced inhibition in its primary input nucleus, the striatum (Mink, 2001; Albin and Mink, 2006; Kurvits et al., 2020). Animal models mimicking this disinhibition may be generated by microinjection or infusion of GABA_A_ antagonists such as bicuculline or picrotoxin into the motor parts of the striatum, which lead to the expression of motor tics phasically over tens of minutes (Tarsy et al., 1978; Crossman et al., 1988; McCairn et al., 2009; Worbe et al., 2009; Bronfeld et al., 2013b; Pogorelov et al., 2015) or tonically over days or weeks (Vinner et al., 2017). This experimental model demonstrates the key symptom of tic disorders, tic expression, however, it is important to note that it does not present the full properties of these disorders such as the premonitory urge or the neurodevelopmental properties (Bronfeld et al., 2013a; Yael et al., 2016). The similarity of tic expression in the experimental model led us to the behavioral hypothesis that the tics would undergo the same modulation observed in TS patients during the wake-sleep cycle. In the experimental model, the timing of a single tic and the body part in which it will be expressed are modulated via the cortical input (Israelashvili and Bar-Gad, 2015) and striatal somatotopic organization (Bronfeld et al., 2013b), respectively, thus enabling a controlled study of tic modulation. The BG have also been implicated in sleep-wake regulation (Vetrivelan et al., 2010; Lazarus et al., 2012, 2013). Neurotoxic lesions (Qiu et al., 2010) and electrostimulations (Qiu et al., 2016) of different nuclei of the BG led to changes in sleep structure. This control in conjunction with the striatal generation of “tic related” activity even without the behavioral expression of tics (Muramatsu et al., 1990) led to the neurophysiological hypothesis that tic related patterns will be generated but will not propagate through the cortico-basal ganglia pathway during sleep.

In this study, we use the chronic experimental model of motor tics in freely behaving rats to quantitatively examine the changes in tic expression during sleep and the underlying changes that occur in the striatum during the sleep-wake cycle. The study of tics during sleep may shed light on the overall mechanism underlying tic expression and its complex modulation by different brain structures in response to internal and external factors.

## Results

### Motor tic expression is reduced during sleep

The GABA_A_ antagonist bicuculline was infused continuously at a fixed rate into the motor parts of the striatum of freely behaving rats, over a period of 7-14 days. During the infusion period, ongoing motor tics appeared in the rats’ head, jaw and/or contralateral forelimb, while the rats expressed the entire naïve behavioral repertoire. This model of chronic motor tic expression in rats was described and characterized in detail in our previous study (Vinner et al., 2017). We conducted 26 recording sessions of a single sleep-wake cycle, in 11 rats during the bicuculline infusion period. The rats’ behavior was monitored using a video stream, and offline manually classified into six behavioral states: quiet waking, sniffing, exploration, grooming, feeding, and sleeping. In this study, we compared the modulation of tics in two behavioral states: the quiet waking state and sleep. The rat was defined as being in a quiet waking state when it was awake, sitting, standing or lying without any additional movement. Sleep was classified as periods when the rat was immobile in a sleeping position with eyes closed or half-closed (van Betteray et al., 1991). These two states of behavior were chosen because of their similarity in terms of the absence of voluntary movement, and their key difference in terms of the vigilance mode. The detailed movement of the rat was monitored using 3D accelerometer, gyroscope and magnetometer motion sensors attached to the head of seven rats across 18 sessions. The X-axis of the gyroscope signal was used for the identification and quantification of tic expression. Throughout the quiet waking state, motor tics manifested in a stereotypic pattern of movement (Fig. 1A), which was distinct from non-tic movements (Fig. 1B). Tic kinematics were stable in terms of shape (Fig. 1C, correlation coefficient across sessions: 0.86 ± 0.14, mean ± SD) and amplitude (Fig. 1E, coefficient of variation (CV) of tic amplitude: 0.45 ± 0.2, mean ± SD) during the quiet waking state. During sleep, the tic frequency decreased significantly compared to the quiet waking state (Fig. 1D, quiet waking, 65 ± 16, sleep 3 ± 6 tics/minute, mean ± SD, t_(17)_=13.76, p<0.001, paired t-test). The tics which did appear during sleep, in 7/18 sessions, were stable throughout the period (Fig. 1E, CV of tic amplitude: 0.49 ± 0.25, mean ± SD; correlation coefficient within the sleep state: 0.84 ± 0.17, mean ± SD). The tics during sleep, however, had significantly lower amplitude compared to the quiet waking state (Fig. 1F, quiet waking 119 ± 73, sleep 5 ± 9 peak-to-peak °/s, t_(17)_=7.01, p<0.001, paired t-test) and did not preserve their typical shape compared to the quiet waking state (Fig. 1G, Correlation coefficient with the quiet waking state: 0.002). The kinematic sensor recordings were employed to characterize and quantify the transition period that was observed during each session. This period covered a continuum from falling asleep that began with the eyes closed or almost closed, contained short waking segments, and terminated with continuous sleep (Fig. 1A) (Lo et al., 2004; Simasko and Mukherjee, 2009). During the transition period, tics continued to appear as frequently as in the quiet waking state (Fig. 1D, 62 ± 17 tics/minutes, mean ± SD, t_(17)_=0.65, p=0.53 and t_(17)_=13.76, p<0.001, paired t-test between the transition period and the quiet waking and sleep states respectively), and preserved their typical shape (Fig. 1C, correlation coefficient within the transition period: 0.83 ± 0.2, median ± SD; Fig.1G, correlation coefficient with the quiet waking state: 0.57), but their amplitude decreased over time (Fig 1C&E, CV of tic amplitude: 0.84 ± 0.33, mean ± SD) and in comparison to the quiet waking state (Fig. 1F, transition period 40 ± 27 peak-to-peak °/s, t_(17)_=5.24, p<0.001 and t_(17)_=5.69, p<0.001, paired t-test between the transition period and the quiet waking and sleep states respectively). Despite the constant infusion of bicuculline, the tic properties changed as a function of the rats’ state. This may suggest the existence of a neural mechanism that bypasses or overcomes striatal disinhibition during sleep.

**Figure 1:**
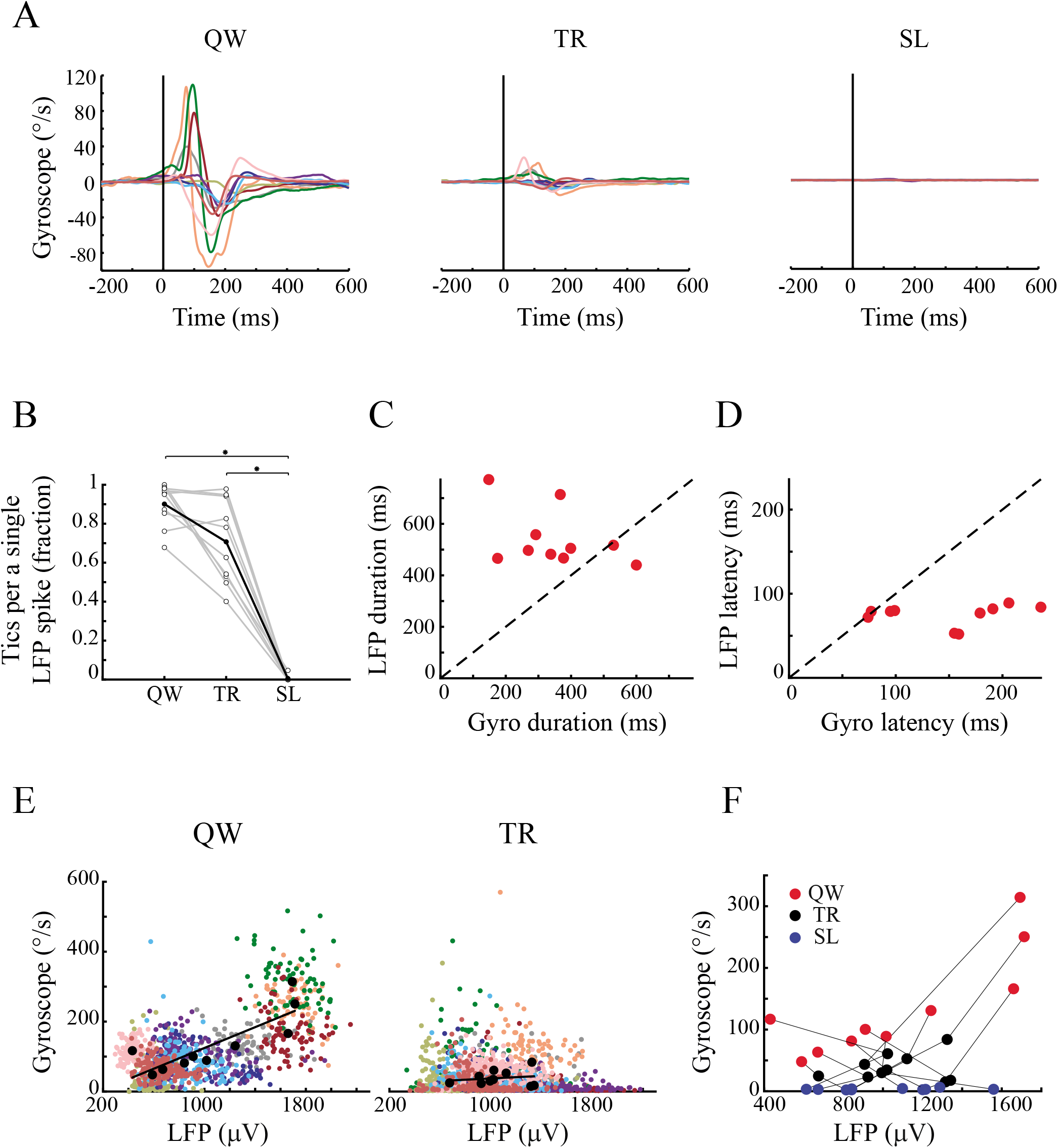
Motor tic frequency and intensity are reduced during sleep. (A) An example of tic expression during the quiet waking state (left), transition period (center) and sleep (right) as recorded in the gyroscope signal, shown using three different time windows. These periods consisted of three distinct behaviors, classified using offline video analysis: quiet waking (red), eyes half-closed (light blue), and eyes closed (blue). (B) An example of video-classified quiet waking (red) and movement (yellow) behaviors as recorded in the gyroscope signal, during ACSF (left) and bicuculline (right) infusions. (C) The peri-tic gyroscope signal (top) and the mean shape for each third (bottom), during the quiet waking state (red shades) and the transition period (shaded black) in a single session (same as A). During sleep, tics were not detected. (D) Tic frequency across all sessions (N=18). Gray lines: individual sessions. Black bold line: mean across all sessions. (E) Coefficient of variation of tic amplitude in the quiet waking state, transition period and sleep. Gray line: mean. (F) Tic intensity, calculated as the peak-to-peak delta across sessions (N=18). Gray lines: single session. Bold black line: mean across all sessions. (G) The mean shape of a motor tic (bold) ± 1 SEM (shadow), across all sessions (N=18) and animals (N=7) in all three behavioral periods (quiet waking - red; transition period - gray; and sleep - blue).

### Macro level neural correlates are dissociated from tic expression during sleep

Extracellular electrophysiological signals in the striatal areas near (<1 mm) the bicuculline infusion site were continuously recorded over 18 sessions in 7 rats, of which 10 sessions from 3 rats included simultaneous kinematic sensor recordings. The neural signal was offline pre-processed to generate three distinct signals: (1) the local field potential (LFP) (frequencies below 100 Hz) which is a summation of activity from multiple sources including synchronous presynaptic input, volume-conducted potentials from remote areas and local potentials (Kajikawa and Schroeder, 2011); (2) Multiunit activity (MUA) (frequencies above 300 Hz) which constitutes the sum of the action potentials from multiple neurons around the recording electrodes (Moran and Bar-Gad, 2010); (3) Single-unit activity, which was extracted from the MUA signal using offline spike sorting to generate spike trains representing individual neurons. During the quiet waking state, within each session, changes in the LFP signal appeared as stereotypic deflections with a typical temporal structure (Fig. 2A-B, correlation coefficient: 0.97 ± 0.02, mean ± SD) and with stable magnitude values (Fig. 2D, CV of magnitude: 0.24 ± 0.10, mean ± SD). These deflections, termed “LFP spikes”, were previously described in the acute tic expression model subsequent to bicuculline injections in primates and rats, and were highly correlated with individual motor tics (McCairn et al., 2009; Israelashvili and Bar-Gad, 2015). Similar results were observed for prolonged periods in our previous study which described the chronic model (Vinner et al., 2017). In the current study, these LFP spikes were also apparent during the transition period (Fig. 2A&B), despite the significant change in tic intensity (as seen in Fig. 1). During sleep, although the observable motor tics declined considerably, the LFP spikes persisted (Fig. 2A) and were stable within sessions in terms of shape (Fig. 2B, correlation coefficient within the sleep state: 0.92 ± 0.11, mean ± SD) and amplitude (Fig. 2D, CV of LFP spike amplitudes: 0.26 ± 0.08, mean ± SD). Differences in LFP spike frequencies and amplitudes were observed within each session between the quiet waking and sleep states (Fig 2B, I&II); however, comparisons across all the neural recording sessions revealed no significant change in the mean LFP spike frequencies (Fig. 2C, quiet waking, 68 ± 20, sleep 60 ± 21 LFP spikes/minute, mean ± SD, t_(17)_=1.81, p=0.09, paired t-test), magnitude (Fig. 2E, quiet waking, 1054 ± 548, sleep 932 ± 366 LFP Spikes/minute, mean ± SD, t_(17)_=1.22, p=0.24, paired t-test) or shapes (Fig. 2F, correlation coefficient between the QW and sleep states: 0.99), in sharp contrast to the changes in the tic kinematics.

**Figure 2:**
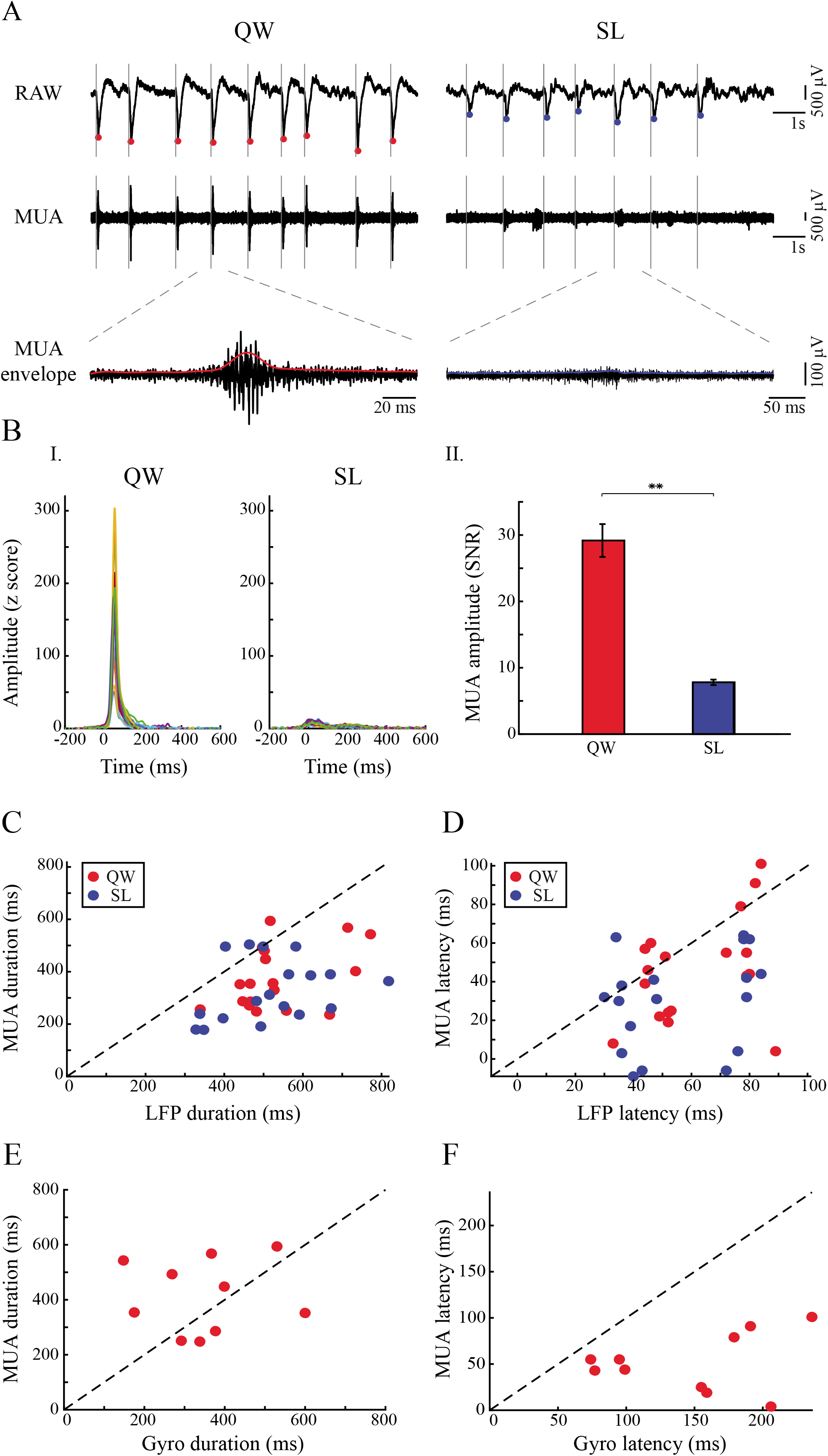
LFP spikes persist during sleep. A) An example of LFP spikes during the quiet waking state (left), transition period (center) and sleep (right) as recorded in the gyroscope signal, in three different time windows. These periods consisted of three distinct behaviors classified using offline video analysis: quiet waking (red), eyes half-closed (light blue), and eyes closed (blue). (B) Peri-LFP spikes (top) and the mean shape for each third (bottom), recorded from the striatum, during the quiet waking state (red shades), transition period (black shades) and sleep (blue shades) in two single sessions (I&II). An example of reduced (I) and increased (II) LFP spikes magnitude during sleep compared to the quiet waking state. (C) LFP spike frequencies across all sessions. Gray lines: single session. Bold black line: mean across sessions. (D) Coefficient of variation of LFP spike amplitudes in the quiet waking and sleep states. Gray line: mean. (E) LFP spike magnitudes across all sessions. The magnitude was calculated as the peak-to-peak delta. Gray lines: single session. Bold black line: mean across all sessions. (F) The mean shape of the LFP spikes (bold) ± 1 SEM (shadow), across all sessions and animals during the quiet waking (red) and sleep (blue) states.

The simultaneous recordings of striatal activity and the kinematic properties enabled the quantification of the relationship between LFP spikes and kinematically-defined motor tics. Multiple analyses demonstrated that while these two signals were highly correlated during the quiet waking state, there was a dissociation between the LFP spikes and motor tics kinematics during sleep: the mean shape of the gyroscope signal around the LFP spikes was the largest in the quiet waking state, decreased in the transition period, and returned to zero during sleep (Fig. 3A). This LFP-movement relation further demonstrates that tics were not simply unidentified due to a lower amplitude during sleep. During the quiet waking state, each detected tic had a high propensity of co-occurrence with a single LFP spike (Fig. 3B, correlation of 90 ± 11%, mean ± SD). Manual examination of the data revealed that even during the low number of cases in which only a motor tic or an LFP spike was identified, but not both, this was due to the low SNR in the second signal which prevented event detection. The dissociation between the events first appeared during the transition period resulting in LFP spikes with no apparent indication of tics. During sleep, almost no motor tics appeared around LFP spike events (Fig. 3A&B, p=0.002 & p=0.002, Wilcoxon signed rank test of sleep compared to quiet waking state and transition states respectively). To study the temporal relationship between the LFP spikes and tics, two parameters were calculated during the quiet waking state for each event: the event duration and the latency to the maximal change. In most of the sessions (8/10) the LFP spike events were longer (542 ± 112 ms, mean ± SD, N=10) than the simultaneous gyroscope deflections (349 ± 141 ms, mean ± SD, N=10) (Fig. 3C, t_(9)_=2.84, p=0.0195, paired t-test). The latency of the LFP spikes (75 ± 13 ms, mean ± SD, N=10) preceded the gyroscope deflection (147 ± 57 ms, mean ± SD, N=10) in 9/10 of the sessions (Fig 3D, t_(9)_=4.02, p=0.003, paired t-test). Further analyses examined the relationship between the magnitude of the LFP spikes and tic intensity. They indicated weak correlations between the magnitude of individual LFP spikes and tic amplitude in 2/10 of the sessions during the quiet waking state, and in 8/10 of the sessions during the transition period (Fig. 3E). Similar analyses, across all the sessions, revealed macro differences between behavioral states, in that the correlation between LFP spike magnitude and the magnitude of the tic was only significant during the quiet waking state (Fig. 3F, *r*^2^ = 0.69, *p* = 0.003).

**Figure 3:**
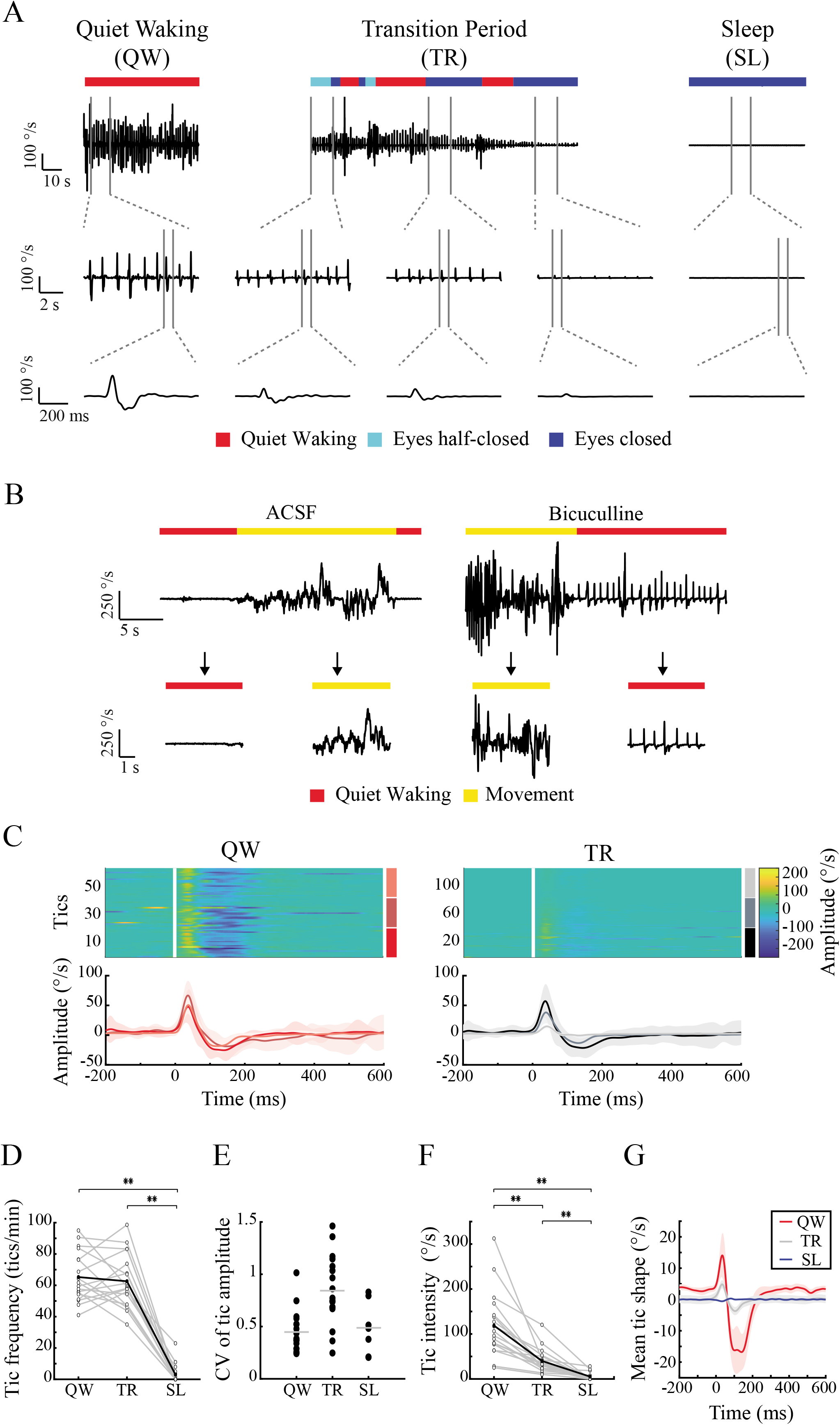
Dissociation of tic expression from LFP spikes during the sleep-wake cycle. (A) The mean shape of the gyroscope signal in each session (different colors), aligned to LFP spike onset. The vertical black line marks the LFP spike onset. (B) The fraction of tic identification around LFP spike events. (C) Comparison of the duration of the mean LFP spike shapes and the related mean gyroscope in each session, during the quiet waking state. (D) Comparison of the latency of the mean LFP spike shapes and the related mean gyroscope deflection in each session, in the quiet waking state. (E) The magnitude of each LFP spike versus the intensity of its subsequent motor tic within sessions (different colors), during the quiet waking state (left) and the transition period (right). Black dots: the mean value in each session, black line: regression line. (F) The mean LFP spike magnitudes in each session (different colors) compared to the mean intensity of the motor tics, during the quiet waking state (red), transition period (gray) and sleep (blue).

### Micro level neural correlates encode tic reduction during sleep

Large changes in the striatal MUA surrounding the injection site were detected in all 191-recorded electrodes. The signal envelope was calculated using a Hilbert transform followed by a low pass filter (Moran and Bar-Gad, 2010) (Fig. 4A). These changes were time-locked to the onset of the LFP spikes and were significantly larger during the quiet waking state than in the sleep state (Fig. 4B, quiet waking, 29.18 ± 2.48, sleep, 7.8 ± 0.41 µV, mean ± SEM, t_(190)_=8.36, p<0.001, paired t-test). The LFP spike-related MUA envelope was assessed using the individual LFP spike onset times as a reference point (peri-LFP spike time histogram - PLTH). The PLTH was calculated using 1 ms bins and was smoothed with a Gaussian window (SD =10 ms). For the PLTH of the MUA envelope, the SNR was calculated by comparing the mean value that preceded (150-200 ms) the tic and its standard deviation. In most of the sessions (17/18 in the quiet waking state and 16/18 in the sleep state), the mean MUA envelope duration (quiet waking, 375 ± 119 ms, mean ± SD; sleep, 327 ± 116 ms, mean ± SD, N=18) was shorter than the mean LFP spike duration (quiet waking-534 ± 114 ms; sleep,518 ± 131 ms mean ± SD, N=18) (Fig. 4C, quiet waking, t_(17)_=5.44, p<0.001, paired t-test; sleep, t_(17)_=5.52, p<0.001, paired t-test). The latency of the MUA (quiet waking, 46 ± 27 ms, mean ± SD; sleep, 30 ± 25 ms, mean ± SD, N=18) preceded the LFP spikes latency in 11/18 sessions during the quiet waking state and in 15/18 sessions during sleep (quiet waking, 62 ± 18 ms, sleep – 56 ± 21 ms,, mean ± SD, N=18) (Fig. 4D, quiet waking, t_(17)_=2.66, p=0.016, paired t-test; sleep, t_(17)_=4.08, p<0.001, paired t-test). There was no significant change between the duration of MUA envelope (414 ± 132 ms, mean ± SD) and the LFP spike-related gyroscope activity (i.e. tics) (349 ± 141 ms, mean ± SD) (Fig 4E, t_(9)_=1.07, p=0.31, paired t-test); however, as expected, the MUA preceded (52 ± 31 ms, mean ± SD) the tic appearance (147 ± 57 ms, mean ± SD) (Fig. 4F, t_(9)_=5.19, p<0.001, paired t-test).

**Figure 4:**
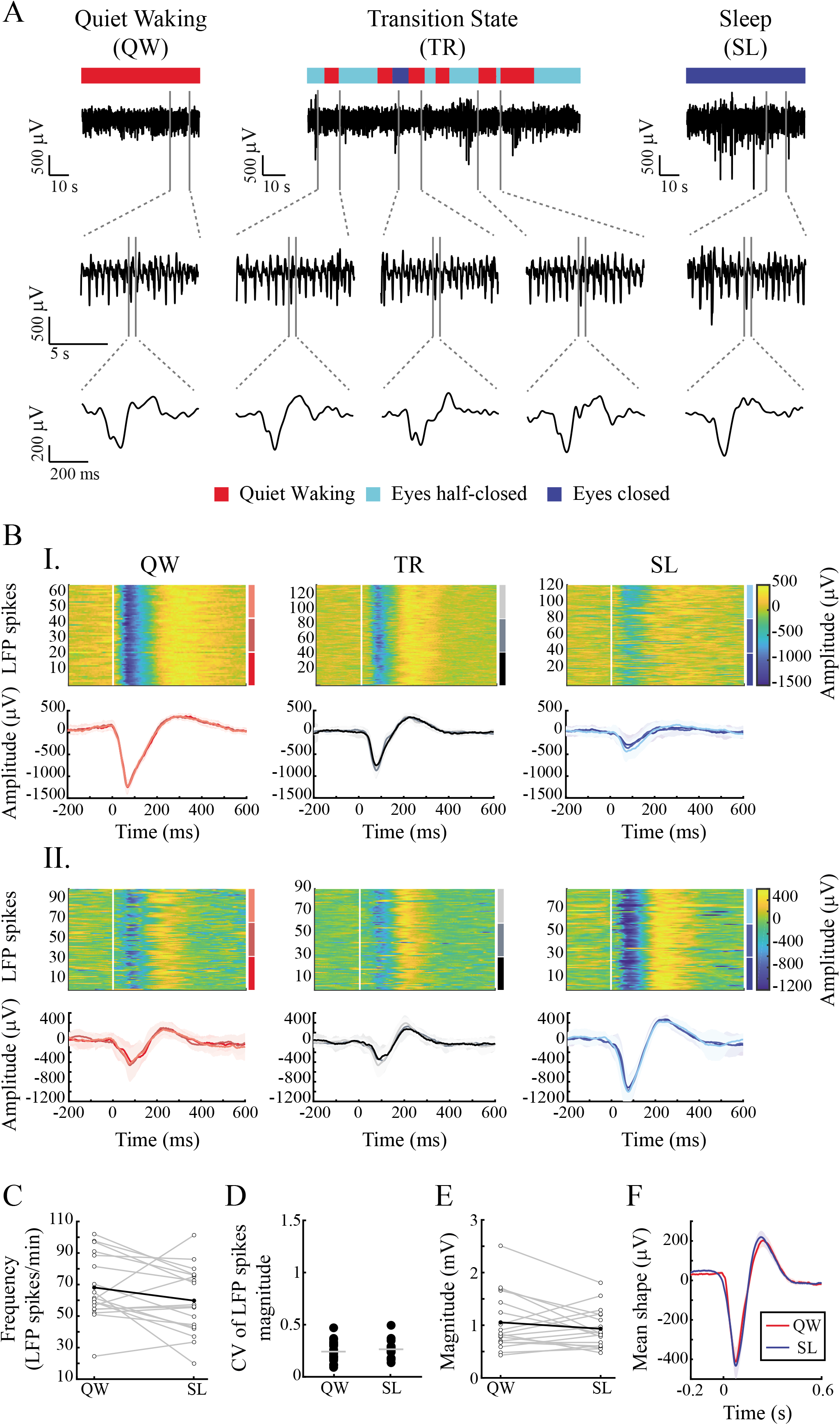
Focal striatal multiunit activity (MUA) decreases during sleep. (A) LFP spike-related MUA (same session as presented in Fig.1A). Left to right: quiet waking versus sleep. Top to bottom: raw signal, striatal MUA, extension of a single MUA episode (black) and its envelop (colored line). Solid gray line: LFP spike onset, dots: LFP spike peaks (quiet waking-red, sleep-blue). (B)(I) An example of a MUA envelope, across all electrodes (different colors) in a single session, aligned to LFP spike onset during the quiet waking and sleep states (same session as presented in Fig.1A, 2A,4A). (II)The mean SNR of LFP spike related MUA envelope ± 1 SEM, calculated from all electrodes (N=191) across sessions (N=18), during the quiet waking and sleep states. (C) Comparison of the duration of the mean LFP spike shapes and the related mean MUA in each session, during the quiet waking state (red) and sleep (blue). (D) Comparison of the latency of the mean LFP spike shapes and the related mean MUA in each session, during the quiet waking state (red) and sleep (blue). (E) Comparison of the latency of the mean MUA envelope and the related mean tic shape in each session, during the quiet waking state. (F) Comparison of the latency of the mean MUA shape and the related mean tic shape in each session, during the quiet waking state.

Neural recordings in the striatum were sorted offline into multiple single-unit spike trains that were then classified into neuronal subtypes based on the spike waveform, firing rate, and pattern. The neurons were identified as SPNs (N=32), and FSIs (N=62). The SPNs firing rate was not affected by the sleep-wake cycle (quiet waking, 2.25 ± 2.02, sleep, 2.30 ± 2.16 spikes/s, median ± SD, p=0.43, Wilcoxon signed rank test), whereas there were significantly lower FSIs firing rates during sleep (quiet waking, 13.12 ± 9.25, sleep – 7.72 ± 6.51, median ± SD, p<0.001, Wilcoxon signed rank test) (Fig. 5A). The firing pattern of both neuron types was more bursty but the change was minor, as evident by the coefficient of variation (CV) (SPNs: quiet waking,1.62 ± 0.4, sleep – 1.77 ± 0.39, median ± SD, p=0.02, Wilcoxon signed rank test, FSIs: quiet waking, 1.31 ± 0.39, sleep, 1.5 ± 0.3, median ± SD, p=0.02, Wilcoxon signed rank test). The LFP spike-related neural activity was assessed with PLTH: A significant response was considered to be excitation or inhibition if it was two SD above or below the mean firing rate of the single cell, respectively. During the quiet waking state, most of the neurons were time-locked (SPNs: 31/32, 97%, FSIs: 58/62 – 94%) to the LFP spike events in different patterns of firing rates. The magnitude of the response of each single cell was calculated as the maximal LFP spike-related firing rate compared to the mean firing rate that preceded (1000-500 ms) the LFP spike. During sleep, most of the neurons were no longer locked in time to the LFP spike events during sleep (SPNs: 12/31 – 39%, FSIs: 28/58, 48%) or exhibited a reduced response that preserved or differed from the firing pattern in the quiet waking state (SPNs: 18/31, 58%, FSIs: 21/58,, 36%) (Fig. 5B). A small portion of the neurons increased their firing rate (SPNs: 1/31, 3%, FSIs: 3/58 similar pattern, 5%, 1/58 different pattern, 2%) or remained unchanged (FSIs: 5/58, 9%). The maximal LFP spike-related firing rate for each single cell demonstrated that in the neural population as a whole this measure was significantly reduced during sleep (Fig. 5C, SPNs: quiet waking, 30.09 ± 41.24, sleep, 6.41 ± 3.86, mean ± SD, t_(31)_=3.29, p=0.002, paired t-test; FSIs: quiet waking, 52.56 ± 41.24, sleep, 19.94 ± 16.61, mean ± SD, t_(61)_=6.07, p<0.001, paired t-test).

**Figure 5:**
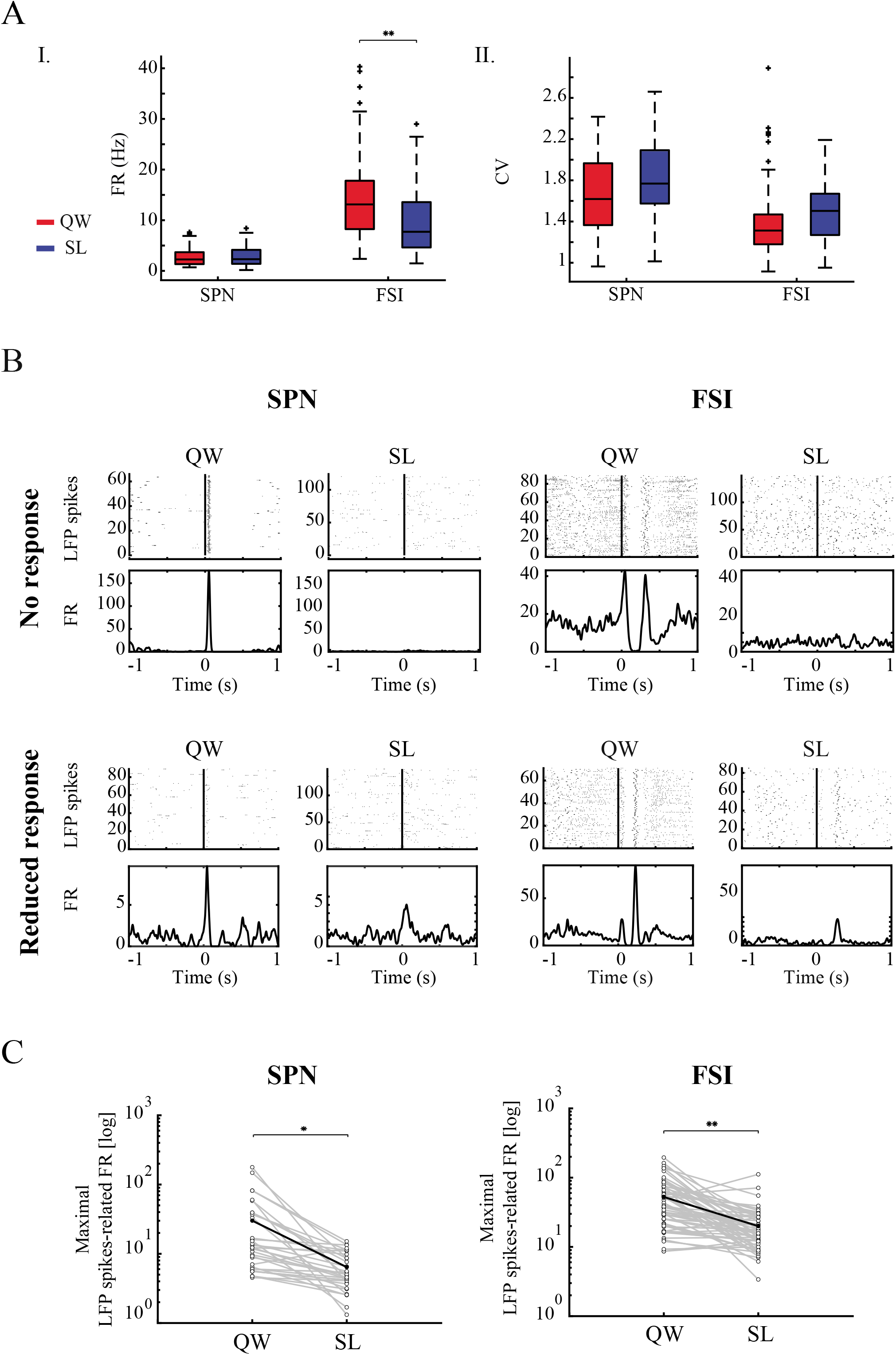
LFP spike related neural activity in the striatum decreased during sleep. (A) I. Mean firing rate for all cell populations. II. Mean coefficient of variation for all cell populations. Quiet waking state (red) and sleep (blue). (B). An example of two SPNs (I), and two FSIs (II), and their different time locks to the LFP spikes during the quiet waking and sleep states. Peri-LFP spike single-neuron raster (top) and peri-LFP spike histogram (bottom). Vertical line: LFP spike onset. (C) The maximal LFP spike-related activity firing rates of all SPNs (left) and FSIs (right) neurons during the quiet waking and sleep states.

## Discussion

We present the first implementation of the chronic model for motor tic expression generated by bicuculline infusion into the striatum via a subcutaneous mini-osmotic pump (Vinner et al., 2017). Throughout the infusion period, the observed tics fluctuated throughout the day depending on the expressed rats’ behavior. We compared two distinct behavioral states: quiet waking and sleep. Both states lack background movements, thus making it possible to compare both behavior and neuronal activity while avoiding distortions generated by excessive movements. The results revealed kinematically stable homogeneous tic expression during the quiet waking state. These tics were significantly reduced in both frequency and intensity during sleep. This behavioral phenomenon was dissociated from the neuronal activity within the striatum: LFP spikes, which are typical of the tic expression state, persisted during sleep despite the absence of observable motor tics. No significant changes in LFP spike shape, frequency or amplitude were observed between states across all sessions despite the large inter-session variance. Multiunit activity near the injection site temporally corresponded to the LFP spike events and was significantly reduced during sleep. Similar LFP spike-related activity occurred at the individual neuron level in striatal neurons belonging to different subpopulations, while presenting diverse changes in firing pattern during sleep.

The effects of sleep on tic modulation in TS patients is still being debated. The earliest reports of Gilles de la Tourette’s patients and the subsequent empirical studies led to the assumption that tics cease during sleep (reviewed by Rothenberger et al., 2001). However, in recent years, evidence from different sources has indicated that tics can appear during sleep in some patients (Champion et al., 1988). Polysomnographic studies have shown that although tics may occur during sleep, they are substantially reduced in both frequency and amplitude (Jankovic and Rohaidy, 1987; Silvestri et al., 1990, 1995; Fish et al., 1991). It has been suggested that tic manifestation during sleep depends on tic severity and may occur during all stages of sleep (reviewed by Oksenberg, 2020). Nocturnal tics were reduced following treatment of the dopamine blocker tetrabenazine (Glaze et al., 1983; Jankovic et al., 1984). Our main results are in line with those observed in TS patients, and demonstrate a significant reduction of tic expression in the experimental model during prolonged sleep. Noticeable tics during sleep were only observed during the transition period, which lasted several minutes and consisted of the falling asleep phase (Wisden and Franks, 2020) intermingled with short awakening segments. We also observed short sequences of awakening during long-lasting sleep episodes (sleep fragmentation) (Press and York, 1980; Trachsel et al., 1991), in which tics reappeared and diminished slowly (as seen in Fig. 6). Reduction in tic intensity was significant while tic frequency was unaffected throughout the transition period, compared to the quiet waking state. These results may imply that tics gradually weakened as the rat fell asleep, such that during sleep, extremely weak tics were not sufficient to cross the observable and kinematical detection threshold. This is further supported by the LFP averaged kinematic signal which demonstrates that even following the averaging process no movement may be detected during sleep.

**Figure 6:**
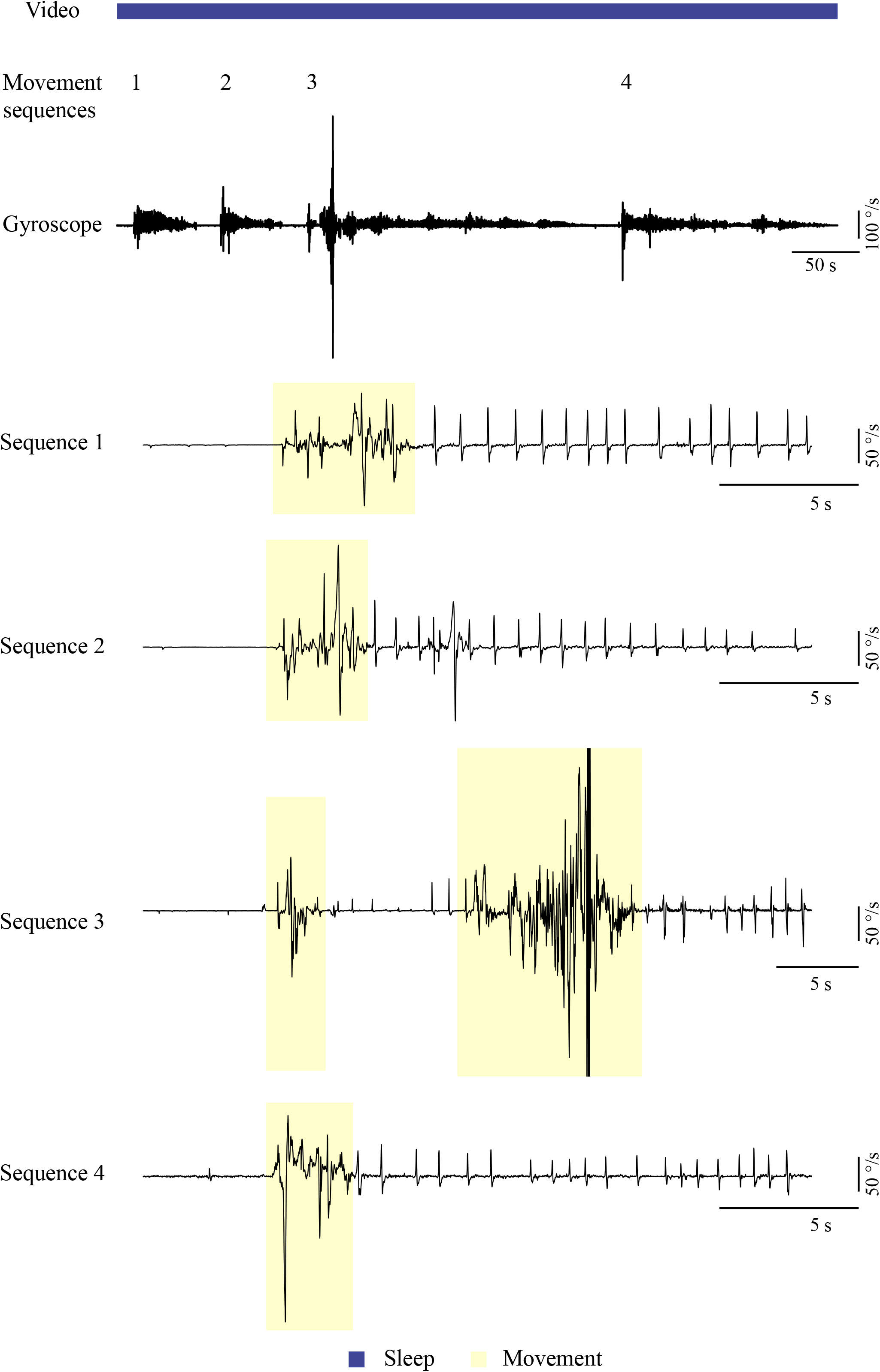
Effect of brief awakenings on tic expression. An example of the gyroscope signal during a long-lasting sleep episode fragmented by short awaking segments. Yellow-movement detection, Blue-sleep state classification.

Sleep is defined by electroencephalographic criteria and divided grossly into two phases: rapid eye movement (REM) sleep and non-REM sleep (Dale Purves et al., 2018). Both phases include excessive skeletal muscle immobility but arise from different neural mechanisms: Non-REM sleep immobility is actively generated by an output nucleus of the BG, the SNr, and indirectly by a chain of processes involving different brain regions (Liu et al., 2020; Wisden and Franks, 2020). REM atonia, on the other hand, is generated by neuronal circuits located in the brainstem that regulate the spinal cord motor neurons (Peever and Fuller, 2017), which is also controlled by BG output (Takakusaki et al., 2004). Tics may occur during different stages of sleep (Jankovic et al., 1984; Jankovic and Rohaidy, 1987; Silvestri et al., 1995; Cohrs et al., 2001) but the evidence for the modulation of their expression during sleep stages is inconsistent (Fish et al., 1991; Silvestri et al., 1995; Cohrs et al., 2001). Sleep attenuates motor symptoms in multiple other movement disorders as well: For example, in Parkinson’s disease, resting tremor diminishes in NREM sleep but may return during awakening and microarousal (reviewed by Garcia-Borreguero et al., 2003) as observed in our study. The disappearance of motor symptoms during sleep has also been documented in animal models of Parkinson’s disease (Buonamici et al., 1986; Degryse and Colpaert, 1986). In our model, we could not classify the different stages of sleep according to the electroencephalogram spectral composition because of the persistence of the LFP spikes, which distorted the apparent spectra. However, the behavioral phenomenon was constant throughout sleep, suggesting that it does not depend on the stage of sleep, but rather on the temporal proximity to wakefulness. The reason for the nocturnal appearance or disappearance of motor symptoms in movement disorders which all involve different pathological abnormalities of BG activity, is still an enigma and should be addressed while considering the critical role of BG in sleep-wake cycle regulation (Urbain et al., 2002; Magill et al., 2004; Mizrahi-Kliger et al., 2018). Our results support the notion that the changes in the activity of the basal ganglia during sleep (loss of tic related activity) plays a key role in addition to the REM and non-REM specific mechanism.

The BG have been shown to play a key role in TS pathophysiology (Peterson et al., 2003; Bloch et al., 2005; Kalanithi et al., 2005; Worbe et al., 2015). LFP recordings performed during deep brain stimulation (DBS) implantation surgeries in TS patients have revealed tonic and phasic changes in BG output associated with tic expression (Zhuang et al., 2009; Israelashvili et al., 2017; Zhu et al., 2019). Extracellular recordings in animal models inducing motor tics have led to a better understanding of the relationship between the properties of neural correlates and tic expression throughout the cortico-BG pathway (Muramatsu et al., 1990; McCairn et al., 2009, 2016; Israelashvili and Bar-Gad, 2015; Klaus and Plenz, 2016). Macro-scale neural correlates, which manifest as deflections of the LFP signal (“LFP spikes”), were shown to be highly correlated with the expression of individual motor tics. The LFP spike magnitude was reported to be the largest in the motor cortex compared to deep structures (McCairn et al., 2009), although significant related neural activity was displayed throughout different BG nuclei simultaneously (McCairn et al., 2009; Israelashvili and Bar-Gad, 2015). Given the shortcomings of the acute model, only awake animals have been studied. Here, using the chronic model, we could reveal a dissociation between these LFP spikes and the expression of motor tics during the spontaneous sleep-wake cycle. LFP spikes were persistent in the striatum during sleep despite the absence of tics. In contrast, LFP spike-related MUA and SUA were significantly reduced during sleep. The dissociation between LFP spikes and tic expression was first documented by Muramatsu and colleagues, where striatal LFP spikes remained present after the cooling or lesioning of the cortex, whereas motor tics were not detectible in the EMG signal (Muramatsu et al., 1990; McCairn et al., 2016). This evidence leads to the assumption that the both LFP spikes and cortical input to the striatum have a role in tic expression. A similar dissociation between LFP spikes and a behavioral expression was also documented in our lab model of hyperactivity, in which striatal disinhibition was limited to the limbic region (Yael et al., 2019). In this case, the abnormal behavior involved overall increased locomotion, whereas the LFP spikes were partially correlated with movement stops. The common mechanism underlying the behavioral changes reflected as motor tics and hyperactivity in these models is believed to involve extensive local activity entrained to the LFP spikes, which is smaller in the limbic regions even though the LFP spikes are dominant (Israelashvili et al., 2020). These results may suggest that the LFP spikes, which are generated due to abnormal and focal disinhibition in the striatum, encode tic generation. The LFP spikes are mandatory but not sufficient for tic expression, but rather the time locking of local networks of neurons in the motor parts of the striatum (reflected in the MUA) is required for the symptom expression. In line with findings, we have previously shown that under the effect of bicuculline injection to the striatum and LFP spikes generation, tic timing is driven by the cortico-striatal activation (Israelashvili and Bar-Gad, 2015). The changes of cortical input to the striatum during sleep affects SPNs’ intracellular properties; from disorganized synaptic events to major shifts in membrane potential (Mahon et al., 2006). The striatum, however, decorrelates its input during sleep (Mizrahi-Kliger et al., 2018), leading to desynchronized output. These changes may thus lead to the reduction of LFP-spike related local neural activity, which in turn may facilitate the disconnection of the tic generation process from the tic expression. The origin of LFP spikes themselves is still an enigma; the origins of LFP signals are diverse and include local synaptic activity, remote synchronized synaptic activity, membrane oscillations, spike hyperpolarization and other sources (Moran and Bar-Gad, 2010). While the local application of bicuculline resulted in a significant MUA and SUA entrained to the LFP spikes implying that the striatum has a key role in LFP spikes generation, the reduction of LFP spikes related neural activity in the striatum during sleep and the unchanged firing rate in the striatum imply that no tic related activity from the striatum affects downstream targets. These results may suggest that other, yet unidentified, brain areas in the CBG loop are the source of LFP spikes.

The current study enables a better understanding of the temporal coding of tics regardless of the behavioral state of the rat. Tic onset time was extracted from the kinematic measures recorded via a wireless system located on the rat’s head. Given the wide spatial distribution of tics, similar to the one occurring in TS patients (Ganos et al., 2015), tics often appeared, separately or simultaneously, in several locations including the rat’s jaw, head, or/and forelimb (Bronfeld et al., 2013b). Therefore, the kinematic properties measured here may have resulted from secondary indirect movement echoing the primary tic. We minimized this effect by analyzing the gyroscope signal and striatal neural activity at onset of the identified LFP spikes which were stable, and independent from the absolute timing of the movement response. We found that the maximal striatal neural (MUA) activity preceded the maximal change of the LFP spikes, and that the tic kinematic peak occurred last. These results are compatible with previous work, in which striatal activity preceded the activity of M1 neurons and were thus considered as a tic initiation reference (Bronfeld et al., 2011) thus underscoring the importance of their role in motor tic encoding.

This work demonstrates a dissociation between tic generation and tic expression, which occur spontaneously and reversibly during the sleep-wake cycle. The coordinated neuronal activity expressed as the LFP spikes encodes the tic generation, which has the potential for expression. However, the behavioral expression of tics depends on the occurrence of extensive striatal local neural activity entrained to the LFP spikes, which subsequently leads to the propagation of this activity to downstream targets. These findings highlight a potential mechanism for tic reduction in TS patients during sleep and potentially their modulation during other behavioral states.

### Limitations of the Study

- The identification of tics in this study is based on kinematic sensors and video and does not include electromyography (EMG) recordings form the specific activated muscles. Thus, we cannot preclude the existence of very low-amplitude tics during sleep. In this case there is still a dissociation of the tic generation from the expression with the change from no-expression to minimal amplitude expression despite equal tic generation.
- The exact brain regions (or combination of regions) within the cortico-basal ganglia pathway which are responsible for the tic generation and expression cannot be pinpointed as this study is limited to the striatum. Further research including simultaneous recordings from multiple regions within the pathway, and their modulation using optogenetic and other measures, are required for determining the relevant brain areas.

### Resource Availability

- Further information and requests for data and code should be directed to the Lead Contact: Izhar-Bar-Gad.
- This study did not generate new unique reagents.
- The code generated during this study is available at **github**: https://github.com/ibglab/ARTICLES

## Methods

### Animals

Eleven adult rats (Long-Evans, females) weighting 269±19 g (mean ± SD) were used in this study. The rats had access to food and water *ad libitum* and were maintained under controlled temperatures and a 12 h light/dark cycle. All procedures were approved and supervised by the Institutional Animal Care and Use Committee and adhered to the National Institutes of Health Guide for the Care and Use of Laboratory Animals and the Bar-Ilan University Guidelines for the Use and Care of Laboratory Animals in Research. This study was approved by the National Committee for Experiments in Laboratory Animals at the Ministry of Health.

### Surgery

An infusion-cannula (stainless steel 30 AWG tube), curved at 90°, was attached to a filled mini-osmotic pump (ALZET pumps, DURECT Corporation), using a flexible polyethylene catheter-tubing (PE-10) and a tubing adapter (CMA Microdialysis). The mini-osmotic pump (9 animals: ALZET Model 2001, volume 200 µl, release rate 1 µl/h, 7 days; 2 animals: ALZET Model 2002, volume 200µl, release rate 0.5 µl/h, 14 days) and the catheter were filled with artificial cerebrospinal fluid (ACSF containing (in mM): 145 NaCl, 15 HEPES, 2.5 KCl, 2MgCl2, 1.2 CaCl2, PH 7.4 with NaOH) or bicuculline methiodide (Sigma-Aldrich) dissolved in ACSF (final concentration: 1 µg/µl for ALZET model 2001 or 2 µg/µl for ALZET model 2002), and then primed in a saline bath at 37 °C for 4-6 hours prior to the implantation surgery. The surgery included insertion of the pump into a subcutaneous pocket in the rat’s back, and implantation of the infusion-cannula into the dorsolateral striatum (infusion target: AP, 1.0 mm; ML, 2.5 mm; DV, 5 mm) (Paxinos and Watson, 2007) to enable ongoing local micro-infusion (Vinner et al., 2017, 2020).

Custom-made movable bundles of 16/32 Formvar-insulated nichrome microwires (25 µm diameter) (Yael et al., 2013) were implanted targeting different basal ganglia nuclei and cortical areas. Only the recording from electrodes targeting the dorsolateral striatum (AP, 0.25 mm; ML, 2.75 mm; DV, 4 mm) and confirmed histologically as being in that location were included in the dataset used for neuronal analysis (N=7 rats). The other animals were only included in the analysis of the behavior.

### Experimental sessions

Initial implantation of bicuculline-filled pumps (3 animals) led to tic expression that commenced during the first day after surgery. In the case of an initial ACSF infusion (8 animals), the rats had a control period of 5-7 days followed by a bicuculline pump exchange and subsequent tic expression. Each experimental session, lasting 100±48 (mean ± SD) minutes, included at least one full spontaneous sleep-wake cycle. During the experimental sessions, neurophysiological and kinematic data were recorded continuously while the animal was awake, moving freely, or sleeping in the recording chamber. The striatal neurophysiological signals were recorded in seven rats across eighteen sessions. The kinematic signals were also recorded in seven rats during eighteen sessions. Of these, ten sessions, recorded from three rats, included simultaneous recordings of both neurophysiological and kinematic signals. The striatal neurophysiological signals were recorded using either a wired system (wide band-pass filtered 0.5–10,000 Hz four-pole Butterworth filter; sampled at 44 kHz; AlphaLab SnR, Alpha Omega Engineering) or a wireless system (wide band-pass filtered 1 Hz single-pole to 7000 Hz three-pole Bessel filter; sampled at 32 kHz; Deuteron Technologies). During recordings from the wireless system, kinematic signals were recorded concurrently using a nine-parameter movement sensor covering the X, Y, and Z axes of an accelerometer, a gyroscope, and a magnetometer (MPU 9150, InvenSense), and were sampled at 1 kHz. All the recorded signals were synchronized with a video stream (30/60 frames/s; HCW850, Panasonic), which enabled offline manual assessment of the behavior.

### Data preprocessing and analysis

The data recorded during each session were composed of three synchronized signals: video, kinematic sensors, and neural activity. The rats’ behavior was extracted from the video stream and was manually classified offline, frame-by-frame, into six behavioral states: quiet waking, sniffing, exploration, grooming, feeding, and sleeping. Motor tics were extracted semi-automatically from the kinematic data. The X-axis of the gyroscope signal was optimal for motor tic identification (as compared to a human expert viewing the video). To compare the quiet waking and sleep states, we sampled the session to obtain two periods of clean and distinct continuous behavior, which lasted 1-3 minutes each. We defined an additional transition period of behavior which consisted of a continuum from falling asleep that began with eyes closed or half-closed, contained short waking segments and terminated with the continuous sleep period. The dataset analyzed in this manuscript only covers these three types of segments, in each session.

The neurophysiological data were preprocessed offline to extract the local field potential (LFP), the multiunit activity (MUA) envelope, and single-unit spike trains. The LFP was extracted using a low-pass filter (100 Hz 4 pole bidirectional Butterworth filter), and the electrode with the largest signal-to-noise ratio (SNR) was chosen. The MUA was extracted using a band-pass filter (300-6000 Hz 6 pole bidirectional Butterworth filter), and for each electrode, the median of all other electrodes within the same session was subtracted. Then, the MUA envelope was calculated using a Hilbert transform (Moran and Bar-Gad, 2010). The single-unit spike trains were extracted from the band-passed signal using cluster-based spike sorting (Offline Sorter, Plexon). Spike trains presenting unstable waveforms or firing patterns were excluded from the database. The single-spike trains were labeled as one of three neuron types (spiny projection neurons – SPNs, tonically active neurons – TANs and fast spiking interneurons – FSIs) according to their firing rate, firing pattern, and waveform shape. As the number of TANs was small (N=_15_), the analysis of their data is not presented. All the offline analyses were performed using custom-written MATLAB code (V2017B; MathWorks).

Motor tics and LFP spikes were detected separately from the gyroscope signal and LFP signal, respectively. Objective identification was performed using the SPRING search strategy (Sakurai et al., 2007). This method utilizes the Dynamic Time Warping (DTW) distance and automatically segments input data into subsequences that match template data. For each session, we created a basic template of a motor tic or a single LFP spike using the first 10 notable events recognized manually. This basic template was then expanded into two templates, where each was a variant of the base template. Each of the templates was fed into the SPRING algorithm, resulting in a list of subsequence matches within the session. LFP spikes or motor tics were defined as subsequences shared by both templates. The onset times of the events were identified to align and average multiple events within a single session, and to explore correlation with other signals.

Throughout the figures -* and ** denote p<0.01 and p<0.001 respectively.

## Acknowledgments

This study was supported in part by an Israel Science Foundation (ISF) grant (297/18) and a BSF-NSF Collaborative Research in Computational Neuroscience (CRCNS) grant (2016744). The authors acknowledge the dedicated work of Diana Olmayev, Lior Bracha, Shai Shkalim, Shani Entin, and Yael Walldman in identifying rat behavior and their ongoing technical assistance. The authors thank Amit Gross for her preliminary study on the effect of anesthesia on tics that supported the main concepts of the present study.

## Author Contributions

**Esther Vinner Harduf:** Conceptualization, Methodology, Investigation, Writing - Original Draft

**Ayala Matzner:** Software, Formal analysis

**Katya Belelovsky:** Methodology, Resources

**Izhar Bar-Gad:** Conceptualization, Writing - Review & Editing, Supervision, Funding acquisition.

## Abbreviations

BG: basal ganglia;
FSI: Fast spiking interneuron;
LFP: local field potential;
REM: rapid eye movement;
SNR: single-to-noise ratio;
SPN: Spiny projection neuron;
TS: Tourette syndrome

